# A comparison of root and shoot hydraulics, aquaporin expression and leaf gas exchange between two grapevine cultivars reveals differences in hydraulic control mediated by aquaporins

**DOI:** 10.1101/224212

**Authors:** Silvina Dayer, Johannes Daniel Scharwies, Sunita Ramesh, Wendy Sullivan, Franziska Doerflinger, Vinay Pagay, Stephen D Tyerman

## Abstract

Hydraulics of plants that take different strategies of stomatal control under water stress are still relatively poorly understood. Here we explore how root and shoot hydraulics, gas exchange, aquaporin expression and abscisic acid (ABA) concentration in leaf xylem sap ([ABA]_xylem_) may be involved and coordinated. A comparison in responses to mild water stress and ABA application was made between two cultivars of *Vitis vinifera* L. previously classified as isohydric (Grenache) and anisohydric (Syrah). Grenache showed stronger adjustments of leaf, plant, and root hydraulic conductances to decreased soil moisture and a steeper correlation of stomatal conductance (*g_s_*) to [ABA]_xylem_ than Syrah resulting in greater conservation of soil moisture, but not necessarily more isohydric behaviour. Under well-watered conditions, changes in vapour pressure deficit (VPD) had a strong influence on *g_s_* in both cultivars with adjustments of leaf hydraulic conductance. Grenache was more sensitive to decreases in soil water availability compared to Syrah that rather responded to VPD. There were stronger correlations between plant hydraulic parameters and changes in aquaporin gene expression in leaves and roots of Grenache. Overall, the results reinforce the hypothesis that both hydraulic and chemical signals significantly contribute to the differences in water conservation behaviours of the two cultivars.

## INTRODUCTION

To withstand abiotic stresses such as drought, plants have evolved complex adaptive mechanisms that are regulated in a dynamic fashion. There is an interplay between stomatal regulation of transpiration (Chaves *et al.*, 2010) and changes in the hydraulic conductivity of roots (*Lp*_r_; Maurel *et al.* (2010)), and leaves (*K*_leaf_; Sack and Holbrook (2006)). The flux of water vapour through stomata, known as transpiration (*E*), is strongly influenced by vapour pressure deficit (VPD; Aphalo and Jarvis (1991), McAdam and Brodribb (2016)) as well as changes in *K*_leaf_ (Pou *et al.*, 2013), through changes of stomatal guard cell turgor. Changes in guard cell turgor involve complex and still debated mechanisms that are mediated by chemical and/or hydraulic signals (Comstock, 2002). It is well established that under soil and/or atmospheric water deficit (WD, i.e. soil drying or high VPD), roots and shoots synthesize abscisic acid (ABA), a plant growth regulator that is translocated to the leaf via the transpiration stream and reaches the guard cells where it induces stomatal closure (Dodd, 2005, Tardieu & Simonneau, 1998). Even though ABA signalling is seen as the main pathway for stomatal regulation, chemical signals other than ABA have been proposed to contribute significantly (Christmann *et al.*, 2007, Wilkinson *et al.*, 2007) including the recently reported γ-aminobutyric acid (GABA; Mekonnen *et al.* (2016)). Christmann *et al.* (2007) demonstrated in *Arabidopsis thaliana* that changes in turgor pressure of leaf mesophyll cells occurred within minutes of root-induced osmotic stress and elicited activation of ABA biosynthesis and signalling required for stomatal closure. These observations support a role of ABA in stomatal closure but call into question whether it acts as the sole primary-long distance signal of water stress.

Hydraulic mediation of stomatal closure has been observed in studies where large diurnal fluctuations of stomatal conductance (*g*_s_) and leaf water potential (Ψ_leaf_) were observed without substantial changes in the soil water content (Salleo *et al.*, 2000). The co-variation of *g_s_* and Ψ_leaf_ has been interpreted as a mechanism to protect the plant from severe dehydration and consequently, xylem cavitation and loss of hydraulic conductivity (Tyree & Sperry, 1989). Other studies have suggested the presence of hydraulic signals based on positive correlations between *K_leaf_* and *g_s_* at a relatively constant Ψ_leaf_ (Nardini *et al.*, 2001). Evidence for the involvement of a hydraulic root-to-shoot signal has been provided by experiments where wild-type tomato plants were grafted on ABA-deficient roots (Holbrook *et al.*, 2002). Despite the inability of the roots to produce ABA, stomata still showed the wild-type response to WD. Recent studies in grapevine suggested that *g_s_* was regulated to a greater degree by hydraulic rather than chemical signals during the early phases of WD, while ABA seemed to have an additive effect involved in the longterm maintenance of stomatal closure under prolonged WD (Tombesi *et al.*, 2015). According to these studies, the involvement of both hydraulic and chemical signals seems to be a more likely explanation in the regulation of *g_s_* under WD.

Aquaporins (AQPs), cellular membrane-bound water channel proteins and members of the major intrinsic protein (MIP) family, have been shown to play a key role in the transcellular or radial flow of water in both leaves and roots (Chaumont & Tyerman, 2014, Steudle, 2000). AQPs are known to be regulated by cytoplasmic pH, divalent cations, and phosphorylation (Li *et al.*, 2015, Tornroth-Horsefield *et al.*, 2006, Tournaire-Roux *et al.*, 2003). In certain species, the transcellular path is a major contributor to *K*_leaf_ (Prado & Maurel, 2013). Rapid and reversible changes in *K*_leaf_ involving AQPs have been observed under fluctuating environmental conditions such as radiation (Prado *et al.*, 2013), WD (Galmes *et al.*, 2007) and in response to exogenous application of ABA (Pantin *et al.*, 2013, Shatil-Cohen *et al.*, 2011). For example in grapevine, *K*_leaf_ decreased by about 30% under water stress concomitantly with a decrease of expression of some plasma membrane intrinsic protein (PIP) and tonoplast intrinsic protein (TIP) AQP isoforms (Pou *et al.*, 2013). Furthermore, these authors found significant positive correlations between *g_s_*, *K*_leaf_ and leaf AQP expression, suggesting a contribution of AQPs in regulating the flow of water. In *Arabidopsis*, xylem-fed ABA reduced *K*_leaf_ by specifically decreasing the water permeability of vascular bundle sheath cells, putatively through inactivation of PIPs (Shatil-Cohen *et al.*, 2011). In line with that study, Pantin *et al.* (2013) confirmed those observations and proposed a model in which ABA close stomata via its already known chemical effect on guard cells (Tardieu & Simonneau, 1998), but also via an indirect hydraulic action through a decrease in leaf water permeability triggered within vascular tissues (Pantin *et al.*, 2013). According to these findings, the hydraulic signal induced by ABA may be an important component in the mechanisms used by different species to regulate *g_s_* under WD.

In addition to *g_s_* and *K*_leaf_ variations, responses to drought include changes in root hydraulic conductivity (*Lp*_r_). In contrast to the commonly observed reduction in *K*_leaf_, ABA application and WD have usually opposite effects on *Lp*_r_: while water stress reduces *Lp*_r_, most studies have reported increased *Lp*_r_ with ABA (Aroca *et al.*, 2006, Parent *et al.*, 2009, Thompson *et al.*, 2007). The increase in *Lp*_r_ by ABA can be interpreted as a mechanism to improve the water supply to the shoot, helping to maintain the water continuum in the plant under soil or atmospheric WD (Kudoyarova *et al.*, 2011, Pantin *et al.*, 2013). Diurnal changes in *Lp*_r_ have been observed under well-watered conditions concomitantly with changes in shoot transpiration (Vandeleur *et al.*, 2009). In general, these variations correlate with the transcript abundance of root AQPs suggesting that water transport across roots is regulated by AQPs to meet the transpirational demand of the shoots (Laur & Hacke, 2013, Sakurai-Ishikawa *et al.*, 2011, Vandeleur *et al.*, 2014). Accordingly, these studies support the hypothesis of shoot-to-root (chemical and/or hydraulic) signalling via the xylem that regulates *Lp*_r_ in response to *E* and that is modulated by AQPs (Vandeleur *et al.*, 2014). Positive correlations between *Lp*_r_, *g_s_* and *E* have also been observed in grapevines exposed to exogenous ABA applications through the soil suggesting a connection between ABA-mediated root and leaf conductances (DeGaris, 2016).

The hydraulic and chemical (ABA-mediated) mechanisms, described above, that operate between roots and leaves to control *g_s_* are particularly important in understanding the contrasting behaviours reflecting more isohydric or anisohydric strategies of various species and even cultivars (varieties or sub-species) to cope with WD (Pantin *et al.*, 2013). In isohydric plants, Ψ_leaf_ is maintained relatively constant under declining soil moisture availability through a tight regulation of *g_s_* (Tardieu & Simonneau, 1998). This behaviour has been reported to confer an advantage of increased drought tolerance (Schultz, 2003) and is thought to be under hydraulic and chemical (ABA) control (Tardieu & Simonneau, 1998). In contrast, anisohydric plants maintain *g_s_* to prioritize photosynthesis, which is related to a more variable Ψ_leaf_ (Tardieu & Simonneau, 1998). This behaviour has been previously reported to operate under chemical control (Tardieu *et al.*, 1996).

The present study aimed to elucidate how the hydraulic behavior, gas exchange and aquaporin expression in two grapevine cultivars previously classified as near-isohydric (Grenache) and anisohydric (Syrah) (Shultz, 2003) differed in their responses to progressive WD, recovery from WD, and exogenous ABA application to roots. We hypothesized that, under increasing WD or exogenous ABA application, the more isohydric Grenache would decrease *Lp*_r_ concomitantly with *g*_s_ and *K*_leaf_ to maintain homeostasis of Ψ_leaf_ mediated by a down-regulation of leaf and root AQPs. In contrast, the relatively anisohydric Syrah should maintain *Lp*_r_ under increasing WD through an up-regulation of root AQPs in order to maintain homeostasis of *g_s_* and *K*_leaf_ but not Ψ_leaf_.

## MATERIALS AND METHODS

### Experimental site and plant material

The experiments were carried out in 2015 and 2016 at The Plant Accelerator^®^, University of Adelaide, Waite Campus located in Urrbrae (Adelaide), South Australia (34° 58’ 17’’ S, 138° 38’ 23’’ E). One-year-old rootlings of own rooted grapevines (*Vitis vinifera* L.) cvs. Grenache and Syrah were planted in 4.5 L pots containing a mixture of 50 % vermiculite and perlite and 50 % of UC soil mix (61.5 L sand, 38.5 L peat moss, 50 g calcium hydroxide, 90 g calcium carbonate and 100 g Nitrophoska^®^ (12:5:1, N:P:K plus trace elements; Incitec Pivot Fertilisers, Southbank, Vic., Australia) per 100 L at pH 6.8. Plants were grown for two months in a temperature-controlled glasshouse (day/night: approx. 25/20 °C) and irrigated to field capacity every three days from December 21^st^ 2015. The vines were pruned to two shoots 10 days after bud burst (January 10^th^, 2016) and oriented upright during their development using wooden stakes. A liquid soil fertiliser (Megamix 13:10:15 N:P:K plus trace elements; Rutec, Tamworth, Australia) at a concentration of 1.6 mL L^-1^ was applied as required to bring all plants to approximately equal size. The fertiliser was applied weekly for three weeks once the plants had developed the first adult leaves. On the 25^th^ February 2016, all vines were moved from the greenhouse and transferred to a DroughtSpotter (Phenospex, Netherlands) automated gravimetric watering platform where individual pots were automatically weighed continuously (15 min intervals) and watered twice daily (0600 h, 1600 h) based on the plant weight loss by transpiration. All plants were irrigated to their field capacity weights (determined the previous days) daily until the start of the experiment. Day and night temperatures in the DroughtSpotter glasshouse were kept at 25/20 °C, respectively.

### Treatments

Grenache and Syrah vines were used to examine the effects of water deficit (WD) and recovery by rewatering (REC) on stomatal conductance (*g_s_*) and root hydraulic conductivity (*Lp*_r_; normalized to root dry weight). A set of vines were kept as control (well-watered; WW), irrigated to field capacity and weight to replace the amount of water consumed by transpiration daily. Water deficit was imposed by reducing the amount of irrigation until a defined value of leaf maximum daily *g_s_* of approx. 50 mmol H_2_O m^-2^ s^-1^ (Medrano *et al.*, 2002) was reached. After maintaining *g_s_* at about 50 mmol m^-2^ s^-1^ for three days, vines were rehydrated by irrigating the pots to field capacity and recovery from water stress was examined after seven days. An additional treatment consisting of an exogenous application of ABA was simultaneously carried out on a separate set of vines from both cultivars. In this treatment, the vines were root-fed by applying 50 μM of ABA (Valent Biosciences Corporation, Libertyville, IL, USA) daily to the root system concurrently with irrigation. The selected concentration of ABA application was based on prior experiments, which showed that 50 μM of ABA applied to the root system of potted vines is required to have a significant effect on *g_s_* (DeGaris et al., 2015). All the pots were covered with a thick layer of perlite to minimize soil water loss through evaporation.

The night before each measurement day, selected vines from each treatment were moved from the DroughtSpotter glasshouse to an adjacent glasshouse with identical environmental conditions for physiological measurements and tissue sampling for gene expression analysis. The measurement days were: i) well-watered (WW) vines only on the day before the experiment commenced (Day 0); ii) WW and WD vines the day water deficit and ABA treatments achieved a *g_s_* of ~50 mmol H_2_O m^-2^ s^-1^ (Day 5); iii) WW, WD, and ABA vines three days after *g_s_* reached ~50 mmol H_2_O m^-2^ s^-1^ and was held constant in WD and ABA vines (Day 7); and, iv) WW and REC vines seven days after rewatering was performed in REC vines (Day 14). At each time point, three to five WW vines were used as ‘controls’ to compare against the specific treatment(s).

### Physiological measurements

#### Leaf gas exchange

In order to track *g*_s_ and establish the desired stress level in WD and ABA treatments, daily measurements of *g_s_* were performed in all vines on the DroughtSpotter platform on fully-expanded leaves (estimated minimum leaf age: Leaf Plastochron Index, LPI >10) in the basal section of the shoots using a porometer (SC-1, Decagon Devices, Pullman, WA, USA). Measurements were performed every day at mid-morning (10:30-11:30 h) on two leaves per treatment and replicate.

On the specific sampling dates, *g*_s_ and transpiration (*E*) were measured concomitantly with the rest of physiological measurements between 10:00 and 11:00 h. Measurements were performed on two fully-expanded, healthy leaves using an open system infrared gas analyzer (LI-6400XT, LI-COR Biosciences Inc., Lincoln, NE, USA) with a 6 cm^2^ cuvette. An external LED light source (LI-6400-02B) attached to the cuvette was used at a fixed PAR value of 1500 μmol m^-2^ s^-1^ due to the non-saturating light levels in the glasshouse for photosynthesis (approx. 200 μmol m^-2^ s^-1^). After gas exchange measurements were performed, the same leaf was excised to determine Ψ_leaf_.

#### Leaf water potential and sap collection for ABA analysis

Predawn, leaf and stem water potentials were measured on adult, primary leaves of vines on the specific sampling dates. Predawn leaf water potential (Ψ_PD_) was measured before sunrise (04:00-05:00 h), and leaf (Ψ_leaf_) and stem (Ψ_stem_) water potentials around midday (11:00-12:00 h). Stem water potential was measured after leaves had been sealed in aluminum foil and plastic bags for two hours, to allow equilibration of water potentials. One leaf per plant was measured from three to five plants per group using a Scholander-type pressure chamber (PMS Instruments Co, Albany, OR, USA).

After recording leaf water potential values, an overpressure of 0.5 MPa was applied to the encapsulated leaf for xylem sap collection (approx. 35 μL). Sap was collected from the cut surface of the protruding petiole using a micropipette and transferred to a pre-weighed and labelled micro tube before snap freezing in liquid nitrogen. Samples were stored at -80 °C until subsequent analysis of ABA.

#### ABA analysis of xylem sap

ABA concentration in xylem sap samples ([ABA]_xylem_) was analysed as described in (Speirs *et al.*, 2013). Briefly, the volume of each sample was measured using a pipette for normalisation. Each sample was mixed with 30 μL of deuterated standard (Plant Biotechnology Institute, Saskatoon, SK, Canada) containing deuterium-labelled analogues of ABA, phaseic acid (PA), dihydrophaseic acid (DPA) and the glucose ester of ABA (ABA-GE) at a concentration of 100 ng mL^-1^ each. Solids were precipitated in a centrifuge at 12,470 × g for 5 min. From each sample, 20 μL supernatant was transferred to a LC/MS tube and analysed by liquid chromatography/mass spectrometry (Agilent 6410 Triplequadropole LC-MS/MS with Agilent 1200 series HPLC, Agilent Technologies Inc. Santa Clara, USA). A Phenomenex C18(2) column (75 mm × 4.5 mm × 5 μm; Phenomenex, Torrance, CA, USA) was used at 40 °C and samples were eluted with a 15 min linear gradient of 10 % to 90 % acetonitrile. Nanopure water and acetonitrile were both mixed with 0.05% acetic acid. Compounds were identified by retention times and mass/charge ratio.

#### Hydraulic conductance of leaves, plant and roots

Leaf and whole plant hydraulic conductance (*K*_leaf_ and *K*_plant_) were determined using the evaporative flux method (Flexas *et al.*, 2013). This measurement is based on the relationship between the driving leaf transpiration rate (*E*) and the water potential gradient (ΔΨ) when leaf water potential reaches a steady state. In this case hydraulic conductance is calculated as follows: *K*_leaf_=*E*/(Ψ_stem_-Ψ_leaf_) and *K*_plant_=*E*/(Ψ_PD_ – Ψ_leaf_).

The hydraulic conductance of the entire root system was measured for the same plants using a High Pressure Flow Meter (Dynamax, Houston, TX, USA) as previously described in Vandeleur *et al.* (2009). This is a destructive technique whereby the stem of the vine is cut above the soil surface, covered with filtered deionized water and the stump connected to the High Pressure Flow Meter with a water-tight seal as quickly as possible, typically within 1 min. A transient ramp in pressure with simultaneous recording of flow rate was used to calculate hydraulic conductance, which was normalized by dividing by total root dry weight to obtain root hydraulic conductivity (*Lp*_r_). All measurements were conducted with 5 minutes of shoot excision. The soil was washed from the roots before drying at 60 °C for more than 48 h prior to weighing.

#### AQPs expression in roots and leaves

Samples of leaves and roots were collected from vines immediately after physiological measurements for subsequent analysis of AQP transcript abundance by quantitative reverse transcription PCR (RT-qPCR). Leaves were immediately immersed in liquid nitrogen and stored at -80 °C until analysis. Roots were carefully selected from the bottom and upper parts of the pot in order to get the thinner, white and more functional roots. The root samples were quickly washed to remove soil particles and dried with tissue paper before being submerged in liquid nitrogen.

Leaf and root material was ground to a fine powder in liquid nitrogen using a mortar and pestle. For leaves, total RNA was extracted from 100 mg of fine frozen powder using the Spectrum Plant Total RNA extraction Kit (Sigma-Aldrich, St. Louis, MO, USA). DNA contamination was avoided by digestion with the On-Column DNase I Digestion Set (Sigma-Aldrich, St. Louis, MO, USA) during RNA extraction according to manufacturer recommendations. For roots, RNA extractions were performed as described by Vandeleur et al. (2014). RNA was extracted from 200 mg of fine frozen powder with a 20 mL sodium perchlorate extraction buffer (5 M sodium perchlorate, 0.2 M Tris pH 8.3, 8.5 % (w/v) polyvinylpolypyrrolidone, 2 % PEG 6000, 1 % (w/v) SDS, 1 % (v/v) β-mercapto-ethanol) for 30 min at room temperature. The lysate was filtered through a glass wool filter and mixed with 30 mL of cold absolute ethanol before precipitation at -20 °C overnight. After centrifugation at 3500 rpm for 20 min at 4 °C, the pellets were washed with cold ethanol and purified using the Spectrum Plant Total RNA Extraction Kit with on-column DNase digestion as described for leaves. Concentration and purity of total RNA were determined on a NanoDrop^™^ 1000 Spectrophotometer (Thermo Fisher Scientific Inc., MA, USA). Agarose gel electrophoresis (1.2 % agarose) was done to visualize the integrity of RNA.

For cDNA synthesis, 1 μg of total RNA was reverse transcribed using iScript^™^ cDNA Synthesis Kit for RT-qPCR (Bio-Rad, CA, USA) according to manufacturer instructions.

Gene expression analysis was carried out by quantitative reverse transcription PCR (RT-PCR) (QuantStudio^™^ 12K Flex; ThermoFisher Scientific Inc., MA, USA). For standard curves, 1:10 serial dilutions of purified PCR products were made in the range of 10^6^ to 10 copies/reaction. RT-qPCR was performed in a 10 μL reaction volume containing 1 μL of either the serial dilution or undiluted cDNA, 5 μL KAPA SYBR^®^ FAST Master Mix (2X) Universal (Kapa Biosystems Inc., MA, USA), 100 nM of gene-specific primers, and 0.2 μL of ROX Reference Dye Low (50X). The thermal cycling conditions were: one cycle of 3 min at 95 °C followed by 40 cycles of 16 s at 95 °C and 20 s at 60 °C. Subsequently, melting curves were recorded from 60 to 95 °C at a ramp rate of 0.05 °C/s. Normalized relative quantities (NRQ) of gene expression were calculated between the gene of interest (goi) and two reference genes *(ELF, GAPDH)* taking differences in PCR efficiency (*E*^Ct^) into account (Hellemans *et al.*, 2007, Pfaffl, 2001):

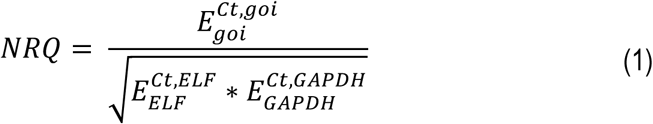

Overall, a mean NRQ value and standard error were calculated from five independent biological replicates. The absence of non-specific products was confirmed by analysis of the melt curves. Sequences for the gene-specific primers were used from Tashiro *et al.* (2016) *(GAPDH)* and Shelden (2008) *(ELF, PIP1;1, PIP2;2, PIP2;3, TIP1;1)* or designed using NCBI Primer-Blast (Ye *et al.*, 2012) for *PIP2;1 and TIP2;1* (Supplementary Table 1). Log_2_ ratios for the heatmap were calculated between the mean NRQ of the WW controls and the mean NRQ of WD, ABA or REC.

#### Statistical Analyses

Analysis of variance (ANOVA) was performed using Infostat software (version 1.5; National University of Córdoba, Córdoba, Argentina). The means were compared using Fisher’s multiple range test (*p* ≤ 0.05) when appropriate, and significant interactions between treatments are indicated and described in the text. Pearson correlation coefficients and multiple linear regressions were calculated in the statistical language R (R Core Team, 2017). Correlation maps were constructed using the R package *corrplot* (Wei and Simko, 2016). Only correlations which were significant at *p* ≤ 0.05 are shown in the correlation maps. Linear regressions and Mann-Whitney test were performed using GraphPad Prism version 7.00 for Windows (GraphPad Software, La Jolla California USA). Figures, except correlation maps, were created in GraphPad Prism.

## RESULTS

### Water relations of mild water deficit and ABA watered vines

Potted vines of Grenache and Syrah were subjected to four different treatments on the DroughtSpotter platform and *g_s_* was monitored using a porometer. Well-watered (WW) vines were maintained at a constant soil water content for maximum *g_s_* (Fig. 1); mild water deficit vines (WD) were not watered until *g*_s_ reached approx. 50 mmol H_2_O m^-2^ s^-1^ (Fig. 1a, Day 4), after which the same deficit was maintained for two more days (Fig. 1b, Day 6); ABA watered (ABA) vines that were watered with 50 μM ABA solution until *g*_s_ reached approx. 50 mmol H_2_O m^-2^ s^-1^ as well (Fig. 1b, Day 6); and, recovery (REC) vines originated from the WD treatment that were re-watered after Day 6 to WW levels (Fig. 1c, Day 13).

**Figure 1.**
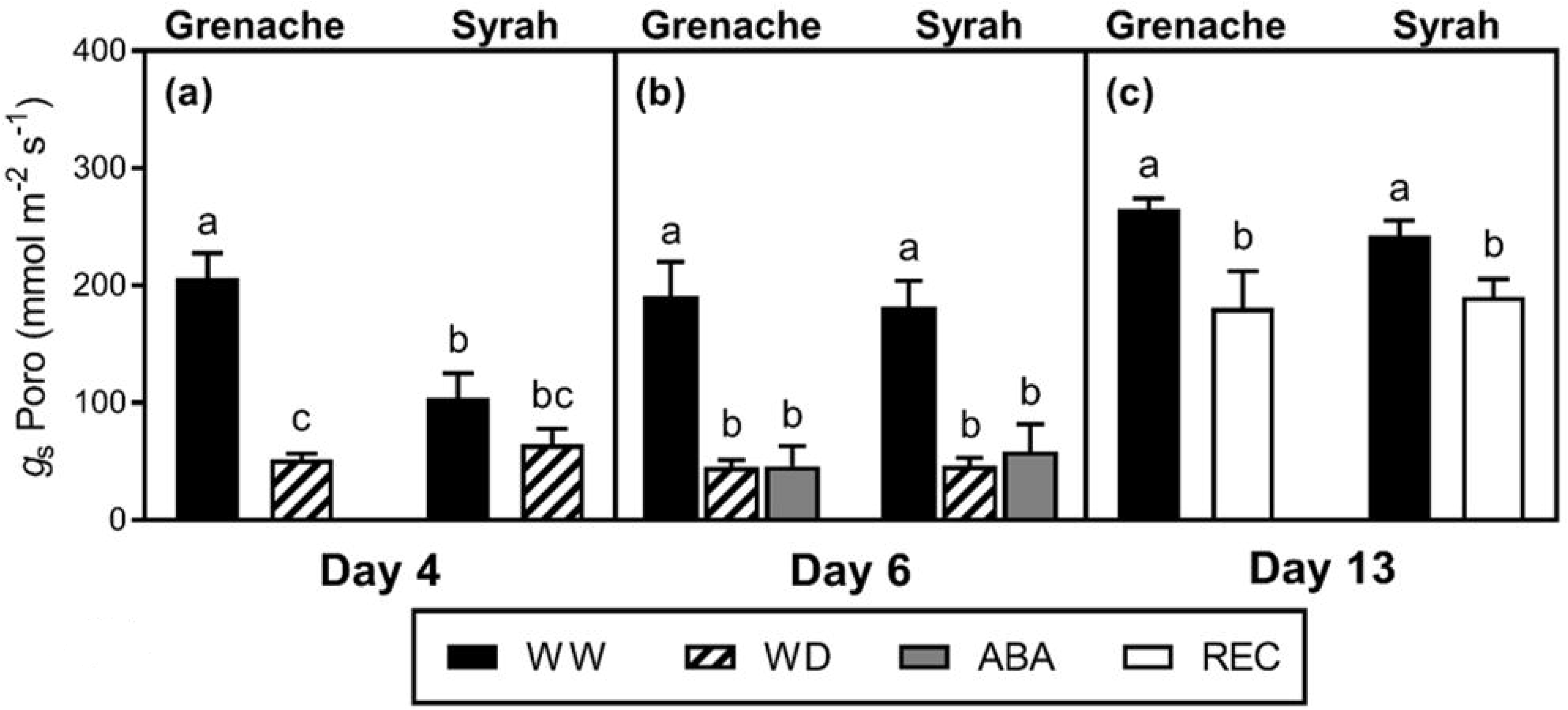
Stomatal conductance as measured with a porometer (g_s_ Poro; a-c) in Grenache and Syrah grapevines under mild water deficit (WD), exogenous application of abscisic acid (ABA) and recovery from water stress (REC) the days before the sampling days along the experiment on the DroughtSpotter platform. Well-watered (WW) vines were measured at each time point and used as control. The WD vines were first sampled at Day 4 when the desired *g_s_* value was achieved (~50 mmol H_2_O m^-2^ s^-1^) and at Day 7 after the stress was sustained for three days. Recovery vines were sampled after seven days of re-watering at WW levels. Values are means ± SE (n = 5). Different letters indicate statistically significant differences between treatments and cultivars (*P ≤ 0.05*) by Fisher’s LSD test.

Due to significant differences in vapour pressure deficit (VPD) between sampling days when physiological measurements were made (Supporting Information Fig. S1), *g_s_* measured by infrared gas analysis (IRGA) varied somewhat, therefore *g_s_* was calculated for the treatments WD, ABA, and REC relative to the WW controls on the same days (Fig. 2a-c). The WD vines of Grenache showed a significantly larger reduction in *g_s_* compared to Syrah on Day 5 (Fig. 2a), while no differences in soil and plant water status (Ψ_PD_, Ψ_leaf_, Ψ_stem_) were observed between WD vines of both cultivars (Fig. 2d,g,j). However, significantly lower Ψ_PD_, Ψ_leaf_, and Ψ_stem_ were measured in WD vines of both cultivars compared to WW vines. A range of Ψ_PD_ which, which is an estimate of soil water potential, from -0.2 to -0.5 MPa was measured in WD vines for days 5 and 7 (Fig. 2d,e). On Day 7, WD vines from both cultivars showed a similar reduction of *g_s_* relative to WW plants after being exposed to the same deficit for 3 days (Fig. 2b), however a significantly lower Ψ_PD_ was measured for WD vines of Syrah compared to Grenache. At the same time, WD vines of Syrah also had a lower Ψ_leaf_ compared to Grenache, while Ψ_stem_ was not significantly different (Fig. 2h,k). The ABA vines, which had a *g_s_* of approximately 50 mmol m^-2^ s^-1^ on Day 6 (Fig.1b) and were not significantly different from the WD vines, showed a significantly larger reduction of *g_s_* relative to the WW and WD vines on Day 7 (Fig. 2b). ABA-treated Grenache vines had significantly lower relative *g_s_* compared to ABA-treated vines of Syrah. Despite adequate soil moisture, similar to the WW vines (Fig. 1e), ABA-treated vines of both cultivars had a similar Ψ_leaf_ and Ψ_stem_ to that of WD vines (Fig. 2h,k). When WD vines were recovered (REC) for seven days, they showed no significant differences in vine water status compared to the WW vines (Fig. 2c,i,l).

**Figure 2.**
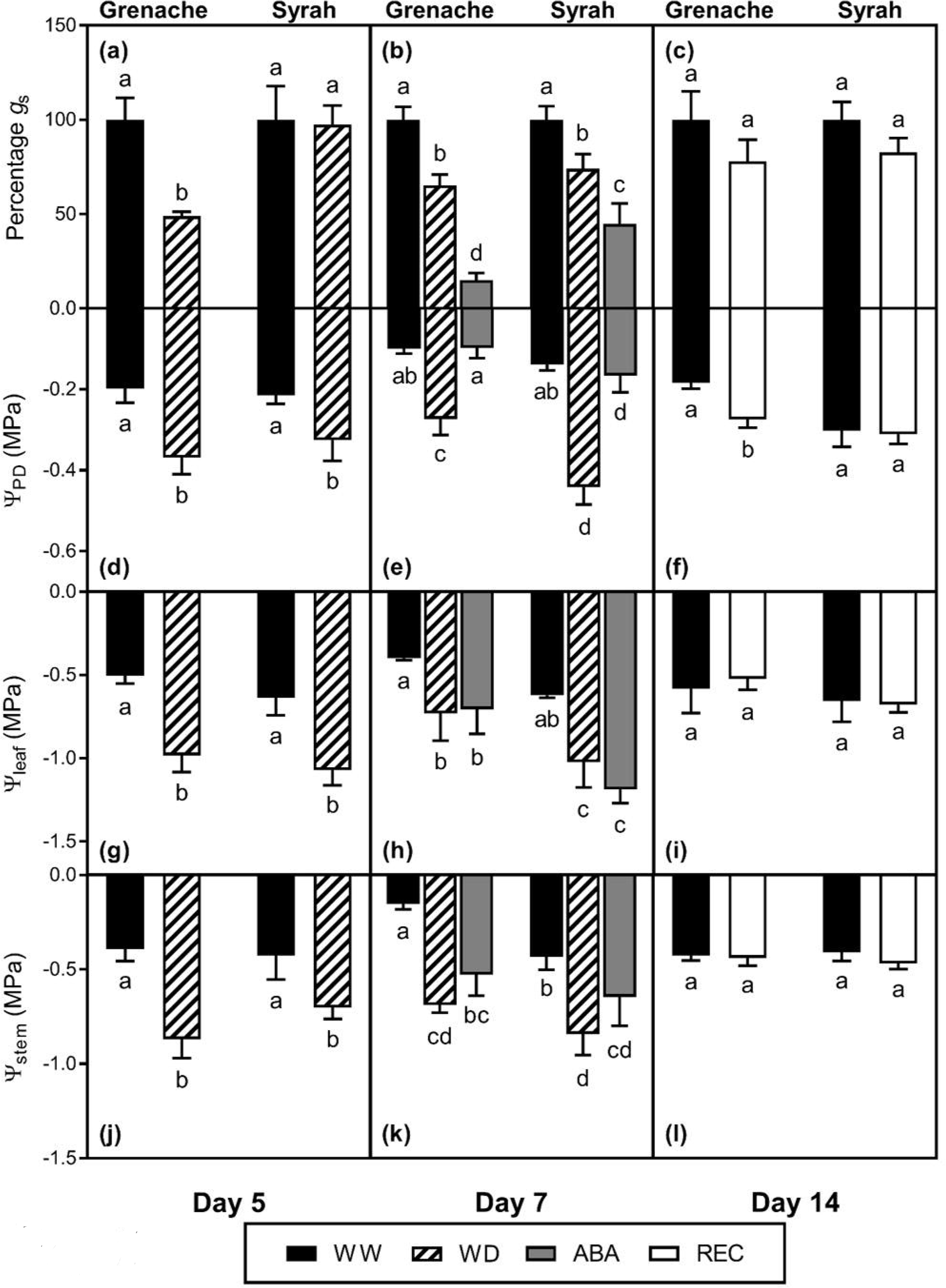
Stomatal conductance relative to WW controls measured with the IRGA (*g*_s_; a-c), pre-dawn (Ψ_PD_; d-f), leaf (Ψ_leaf_; g-i) and stem (Ψ_stem_; j-l) water potentials in Grenache and Syrah grapevines under mild water deficit (WD), exogenous application of abscisic acid (ABA) and recovery from water stress (REC) at different sampling days along the experiment in an adjacent glasshouse to the DroughtSpotter platform. Values are means ± SE (n = 5). Different letters indicate statistically significant differences across all treatments and cultivars within the day (*P ≤ 0.05*) by Fisher’s LSD test.

### Correlations between physiological parameters and aquaporin gene expression

The statistical relationship between the measured plant physiological parameters and gene expression of selected aquaporins is shown in the split correlogram for WW vines in Fig. 3a and for the combination of WW and WD vines in Fig. 3b. The data for Grenache are shown in the upper triangle and data for Syrah are shown in the lower triangle of each correlogram divided by a black diagonal line. Specific categories of columns (col) are labelled along the bottom (A to F) and rows (row) on the right hand side (1 to 6) to enable reference (e.g. col A, row 1) to relevant sets of correlations.

**Figure 3.**
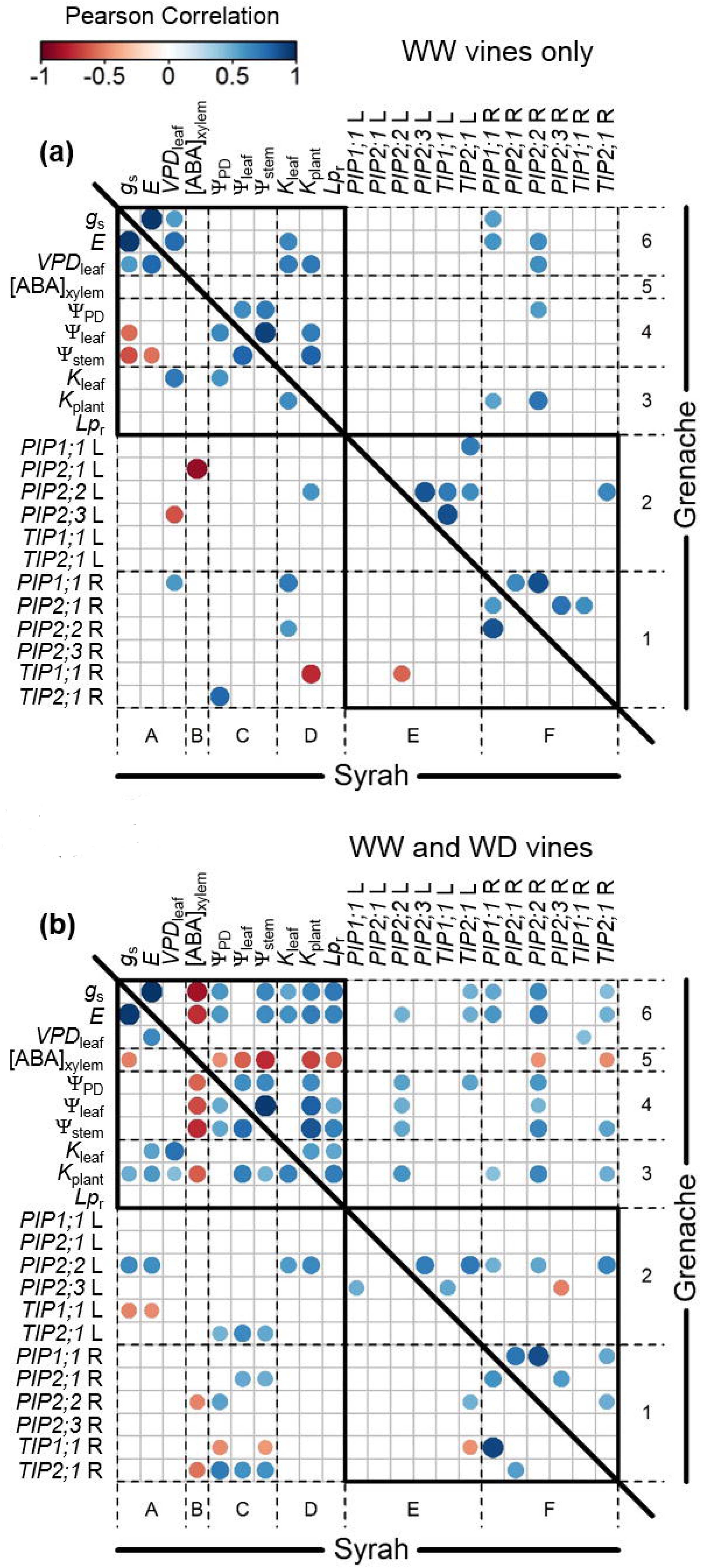
Correlation maps (correlograms) of plant physiological parameters and gene expression in leaves and roots of the cultivars Grenache (upper triangle) and Syrah (lower triangle). (a) Correlations based on data from well-watered vines (WW) only and (b) correlations based on combined data from well-watered (WW) and mild water deficit (WD) vines. The exogenous ABA treated vines and recovery from mild water deficit (REC) treatments were excluded from the analysis. Only significant Pearson correlations coefficients (P ≤ 0.05) are shown. Significant positive correlations are shown in blue, while significant negative correlations are shown in red. [ABA]xylem was log10 transformed before the analysis. Expression of aquaporin genes measured in leaves are denoted with an “L” (e.g. *PIP1;1* L), while expression in roots are denoted with an “R” (e.g. *PIP1;1* R).

Since environmental conditions, like vapour pressure deficit (VPD, Supporting Information Fig. S1) changed significantly between different days of the experiment, measurements on WW vines were also variable. A correlogram for WW vines only (Fig. 3a) was used to explore the primary sources and effects of this variability in the data, by excluding all other treatments. Within the recorded physiological parameters (upper left black box), some were significantly correlated to *VPD*_leaf_. Gas exchange parameters, *g_s_* and *E*, were positively correlated to *VPD*_leaf_ for well-watered vines in both cultivars (col A, row 6, Fig. 3a). Additionally, *K*_leaf_ and *K*_plant_ were positively correlated to *VPD*_leaf_ for well-watered vines of Grenache (col D, row 6, Fig. 3a) but not for *K*_plant_ in Syrah (col A, row 3). No significant correlation was found between root hydraulic conductance (*Lp*_r_), and *VPD*_leaf_. Also, expression of some AQP genes was correlated to *VPD*_leaf_. Gene expression of *PIP2;2* in roots was positively correlated to *VPD*_leaf_ in well-watered vines of Grenache (col F, row 6), while in well-watered vines of Syrah, *PIP2;3* in leaves and *PIP1;1* in roots were negatively and positively correlated to *VPD*_leaf_, respectively (col A, rows 2,1). While *g*s and *E* increased with increasing *VPD*_leaf_ for both cultivars, only Syrah showed negative correlations with Ψ_leaf_ and Ψ_stem_ (col A, row 4; Fig. 3a). ABA concentration in the xylem of leaves from well-watered plants ([ABA]_xylem_) showed very few significant correlations to other parameters in general (col B and row 5, Fig. 3a). None of the gas exchange and hydraulic conductance parameters correlated to xylem ABA concentration [ABA]_Xylem_, however [ABA]_xylem_ of WW vines from Grenache (143.5 ± 25.26 ng mL^-1^, n = 13) was higher (P = 0.0019) compared to leaves of WW vines from Syrah (43.77 ± 3.731 ng mL^-1^, n = 11).

Data from WW and WD vines were combined and a second correlogram was developed to investigate the relationships between plant physiological parameters and root and leaf AQP gene expression during the transition from well-watered to water deficit conditions (Fig. 3b). In contrast to WW vines only, *g*_s_ was not significantly correlated to *VPD*_leaf_ for both cultivars once data of WD vines were included, but negative correlations were observed to [ABA]_xylem_ (col A, row 5 and col B, row 6; Fig. 3b). This correlation was stronger in Grenache where *g*_s_ also correlated with Ψ_PD_ and Ψ_stem_ (col C, row 6, Fig. 3b) unlike Syrah. **VPD*_leaf_* correlated positively to *E* only in Syrah but not in Grenache. *K*_leaf_ and *K*_plant_ (col D row 6, Fig. 3b) also showed correlations to *g_s_* and *E* in the cultivar Grenache. While *K*_leaf_ was not correlated to [ABA]_xylem_ in both cultivars, *K*_plant_ (*r* = -0.60, *P* = 0.005) was negatively correlated to [ABA]_xylem_ for both cultivars (col D row 5 & col B row 3, Fig. 3b). *Lp*_r_ was positively correlated with *E*, *g_s_*, Ψ_leaf_ and Ψ_stem_ only for Grenache where it was also negatively correlated to [ABA]_xylem_ (col D, rows 3-6).

The expression of AQPs showed different patterns between the two cultivars (compare top right quadrant for Grenache (cols E-F, rows 3-6) with bottom left quadrant for Syrah (cols A-D, rows 1-2, Fig. 3b). Notable differences are: the absence of negative correlations between leaf *TIP1;1* and gas exchange parameters in Grenache compared to Syrah (col A, row 2) and the absence of positive correlations between leaf *TIP2;1* and *g_s_* and *E* in Syrah compared to Grenache (col E row 6). However, leaf *TIP2;1* in Syrah was more positively correlated with Ψ_leaf_ (col C row 2). In Grenache, expression of leaf *PIP2;2* was positively correlated to E, water potentials and *K*_plant_ (col E rows 3-6), while for Syrah positive correlations with *g_s_*, *E*, *K*_leaf_ and *K*_plant_ (cols A-D row 2) were observed.

The two most highly expressed aquaporins in roots *(PIP1;1* and *PIP2;2)* also showed differences between the two cultivars (cols A-D row 1 and col F rows 3-6). Root *PIP1;1* for Grenache was positively correlated with *g_s_*, *E* and *K*_plant_ while Syrah showed no significant correlations for these parameters. Similar to leaves, expression of *PIP2;2* in roots of Grenache was positively correlated to *g_s_* and *E*, water potentials and *K*_plant_ (col F rows 3-6), but not to *Lp*_r_. In Syrah these were notably absent except for a positive correlation with Ψ_PD_. Root *TIP2;1* showed positive correlations to *g_s_*, *E*, Ψ_stem_ and *K*_plant_ in Grenache, but only positive correlations to water potentials in Syrah (similar to leaf *TIP2;1).* In both cultivars, root *PIP2;2* and *TIP2;1* were negatively correlated to [ABA]_xylem_ (col B row 1 and col F row 5). Interestingly, no significant correlations were found between any aquaporins and *Lp*_r_.

### Influence of changes in VPD on well-watered vines

A closer examination of the correlation between *g*_s_ and *VPD*_leaf_ (Fig. 3a) in WW vines of Grenache (*r* = 0.57, *P* = 0.034) and Syrah (*r* = 0.56, *P* = 0.036) revealed a similar response of increasing *g_s_* with increasing VPD in the range of 2 to 3 kPa, but significantly higher *g_s_* in Grenache compared to Syrah (Fig. 4).

**Figure 4.**
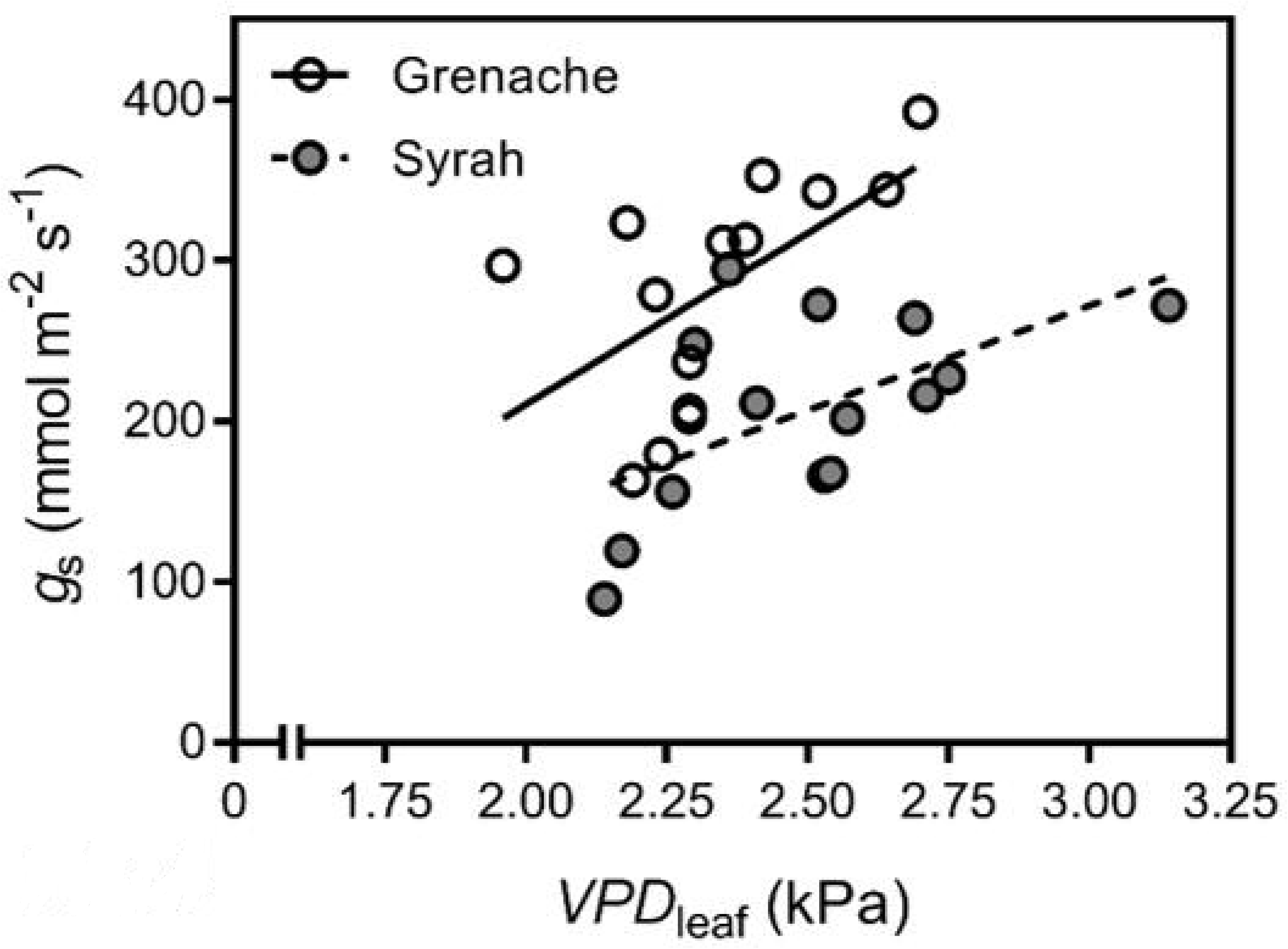
Relationship between stomatal conductance (*g*_s_) and *VPD*_leaf_ for well-watered vines (Days 0, 5, 7, 14) during the experiment (both measured with the IRGA). The slopes for both linear regressions are significantly positive (*P* ≤ 0.05), but not significantly different between the two cultivars. However, the elevations are significantly different (*P* = 0.0002).

While the correlogram for WW vines (Fig. 3a) showed that *K*_leaf_ was only positively correlated to *E* in the cultivar Grenache, linear regressions show that data from both Grenache and Syrah fall on the same line (Fig. 5). The correlation between *K*_leaf_ and *E* is just short of being significantly positive (*P* = 0.052). Grenache had a significantly higher *K*_leaf_ relative to Syrah under well-watered conditions (Fig.5 inset). Similar to *K*_leaf_, significantly higher *K*_plant_ were measured for the cultivar Grenache, compared to Syrah (Gr.: 18.5 ± 2.2 mmol MPa^-1^ m^-2^ s^-1^, n = 14; Sy.: 11.1 ± 1.0 mmol MPa^-1^ m^-2^ s^-1^, n = 14; *P* = 0.0058) and *Lp*_r_ (Gr.: 7.3 x 10^-6^ ± 1.0 x 10^-6^ kg s^-1^ MPa^-1^ g^-1^, n = 11; Sy.: 1.7 x 10^-6^ ± 5.3 x 10^-7^ kg s^-1^ MPa^-1^ g^-1^, n = 12; *P* = 0.0002). Correlations between *E* and *K*_plant_ were not significant at *P* ≤ 0.05, but correlation coefficients were positive for both Grenache (*r* = 0.50, *P* = 0.067), which was close to being significant, and Syrah (*r* = 0.37, *P* = 0.194). The correlations between *E* and *Lp*_r_ were also not significant at *P* ≤ 0.05, but for Grenache (*r* = 0.51, *P* = 0.108) a positive correlation coefficient was found, while for Syrah (*r* = 0.08, *p* = 0.796) the correlation coefficient was close to zero. The water potential gradient between stem and leaf (Ψ_stem_ – Ψ_leaf_) did not show any significant change in well-watered vines of both cultivars in relation to changes in *E (*r** < 0.15, *P* > 0.6).

**Figure 5.**
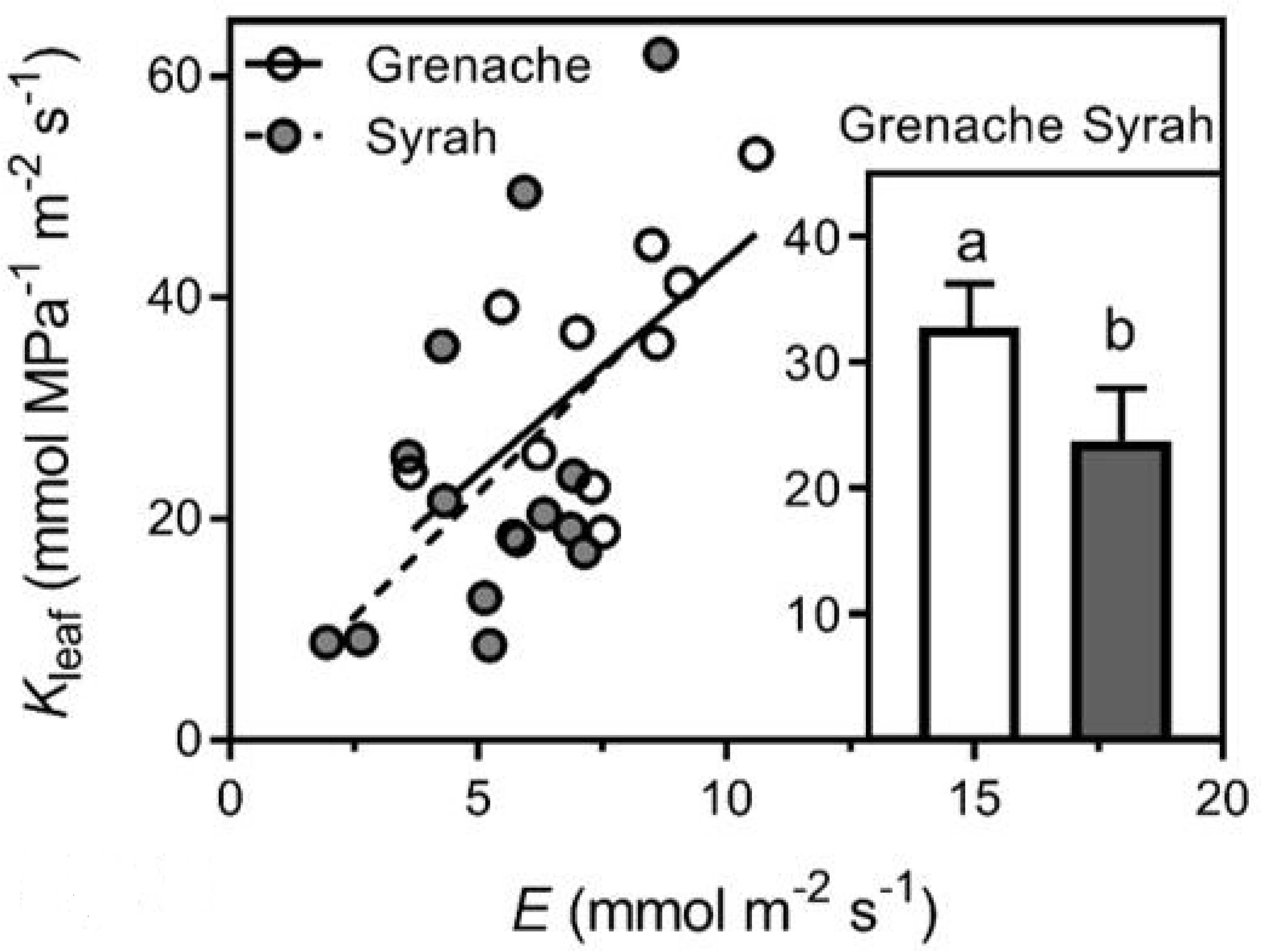
Relationship between *K*_leaf_ and *E* for well-watered vines. For both cultivars, the points fall on the same line. In Grenache, this positive correlation is significant (*r* = 0.65, *P* = 0.030) while in Syrah it is almost significant (*r* = 0.53, *P* = 0.052). Inset: comparing the mean *K*_leaf_ between the two cultivars showed that Grenache had a significantly higher *K*_leaf_ compared to Syrah (*P* = 0.044, Mann-Whitney test).

### Effects of mild water deficit and ABA treatment in Grenache and Syrah

A multiple linear regression model developed for each cultivar indicated that in Grenache, the soil water availability, given by Ψ_PD_is, is the main factor that explains changes in *g_s_* and overrides any effect of changes in *VPD*_leaf_. In contrast, changes in *g_s_* in Syrah are primarily explained by *VPD*_leaf_ (Table 1).

**Table 1.** Multiple linear regression model to predict changes in stomatal conductance (*g_s_*) in response to environmental changes like vapour pressure deficit (*VPD*_leaf_) and predawn water potential (Ψ_PD_), which reflects the soil water availability. Model formula = *g_s_* ~ *VPD*_leaf_ + Ψ_PD_. Input: all data pooled from well-watered, water deficit, and recovery vines from the cultivars Grenache and Syrah. The variables *VPD*_leaf_ and Ψ_PD_ were centred before running the multiple linear regression.

|  | Grenache |  |  | Syrah |  |  |
| --- | --- | --- | --- | --- | --- | --- |
|  | Estimate | Std. Error | P value | Estimate | Std. Error | P value |
| (Intercept) | 223.76 | 15.06 | < 0.001 *** | 177.15 | 10.76 | < 0.001 *** |
| $VPD_{leaf}$ | 16.46 | 74.07 | 0.826 | 85.94 | 40.81 | 0.046 * |
| $\Psi_{PD}$ | 557.36 | 160.06 | 0.002 ** | 84.81 | 107.89 | 0.440 |

In Grenache, *g_s_* significantly decreased (*P* = 0.004) when Ψ_PD_ dropped from approx. -0.1 MPa to -0.5 MPa, while no significant change of *g_s_* was observed in Syrah (Fig. 6a). However, in both cultivars Ψ_leaf_ declined significantly (Grenache: *P* = 0.002; Syrah: *P* = 0.018) in response to decreasing Ψ_PD_ (Fig. 6b). Simple linear regressions showed that the relationship between Ψ_PD_ and Ψ_leaf_ was similar for both cultivars with a slope > 1. Likewise, Ψ_PD_ and Ψ_leaf_ measured on field grown well-watered and droughted Grenache and Syrah by Schultz (2003) also showed the same relationship with a slope of 0.7 (Supporting Information Fig. S2).

**Figure 6.**
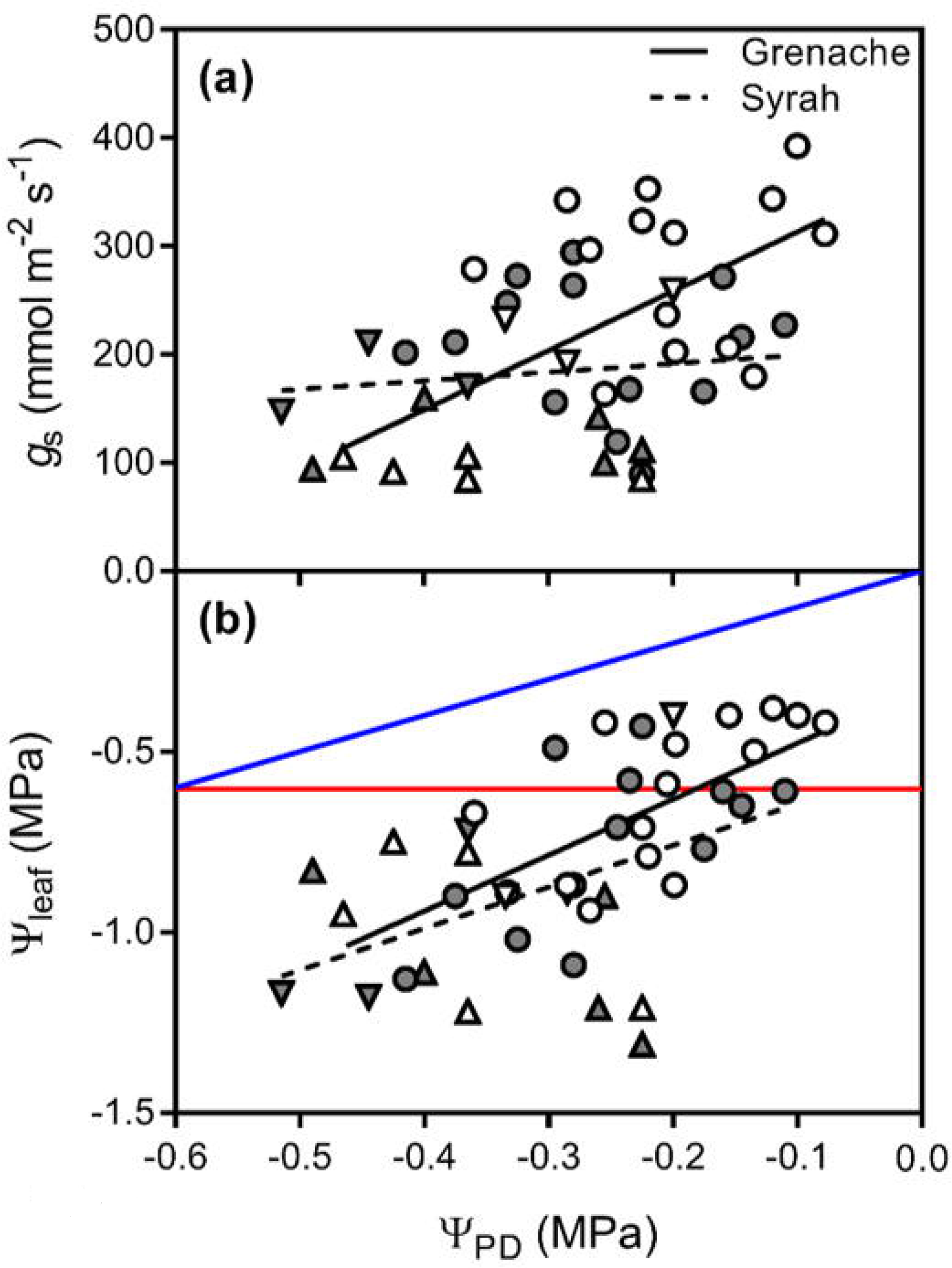
Relationship between Ψ_PD_, *g_s_* and Ψ_leaf_ for well-watered (circles) and water deficit (Day 5: triangles up, Day 7: triangles down) vines. a) Grenache and Syrah show a significantly different (*P* = 0.030) relation between Ψ_PD_ and *g_s_*. While in *g_s_* is significantly positively correlated to Ψ_PD_ in Grenache (*r* = 0.59, *P* = 0.004), no significant correlation was found for Syrah (*r* = 0.12, *P* = 0.536). b) For both cultivars, Ψ_leaf_ is significantly positively correlated to Ψpd (Grenache: *r* = 0.62, *P* = 0.002; Syrah: *r* = 0.50, *P* = 0.018). No significant differences were found for the slope (*P* = 0.531) and intercept (*P* = 0.146) of the linear regression lines (Grenache: Ψ_leaf_ (MPa) = 1.5 ± 0.4 x Ψ_PD_ (MPa) -0.3 ± 0.1; Syrah: Ψ_leaf_ (MPa) = 1.2 ± 0.4 x Ψ_PD_ (MPa) -0.5 ± 0.1). Linear regression lines with a slope of 1 and y-intercept of zero (blue solid line) and slope of 0 and y-intercept of -0.6 (red solid line), which equals the mean Ψ_leaf_ of well-watered Grenache vines, are shown as reference.

The two cultivars were compared for the relationship between *g_s_* and [ABA]_xylem_ in Fig. 7. The decline in *g_s_* with increasing [ABA]xylem for Syrah was significantly (*P* = 0.0005) lower than that for Grenache. Note the logarithmic transformation for [ABA]_xylem_ indicating that the relationship is not linear but exponential. Vines that were treated with ABA instead of water deficit, showed the same relationship between [ABA]_xylem_ of leaves and *g_s_* for both cultivars, and these data are included as squared symbols in Fig. 7. The slopes of the linear regression lines were very similar for data from well-watered and water deficit vines (Grenache: -150.3 ± 26.2 mmol H_2_O m^-2^ s^-1^ ng^-1^ mL; Syrah: -45.6 ± 22.2 mmol MPa^-1^ m^-2^ s^-1^ ng^-1^ mL) and additionally with data from ABA treated vines included (Grenache: -166.4 ± 23.9 mmol H_2_O m^-2^ s^-1^ ng^-1^ mL; Syrah: -53.3 ± 18.3 mmol H_2_O m^-2^ s^-1^ ng^-1^ mL).

**Figure 7.**
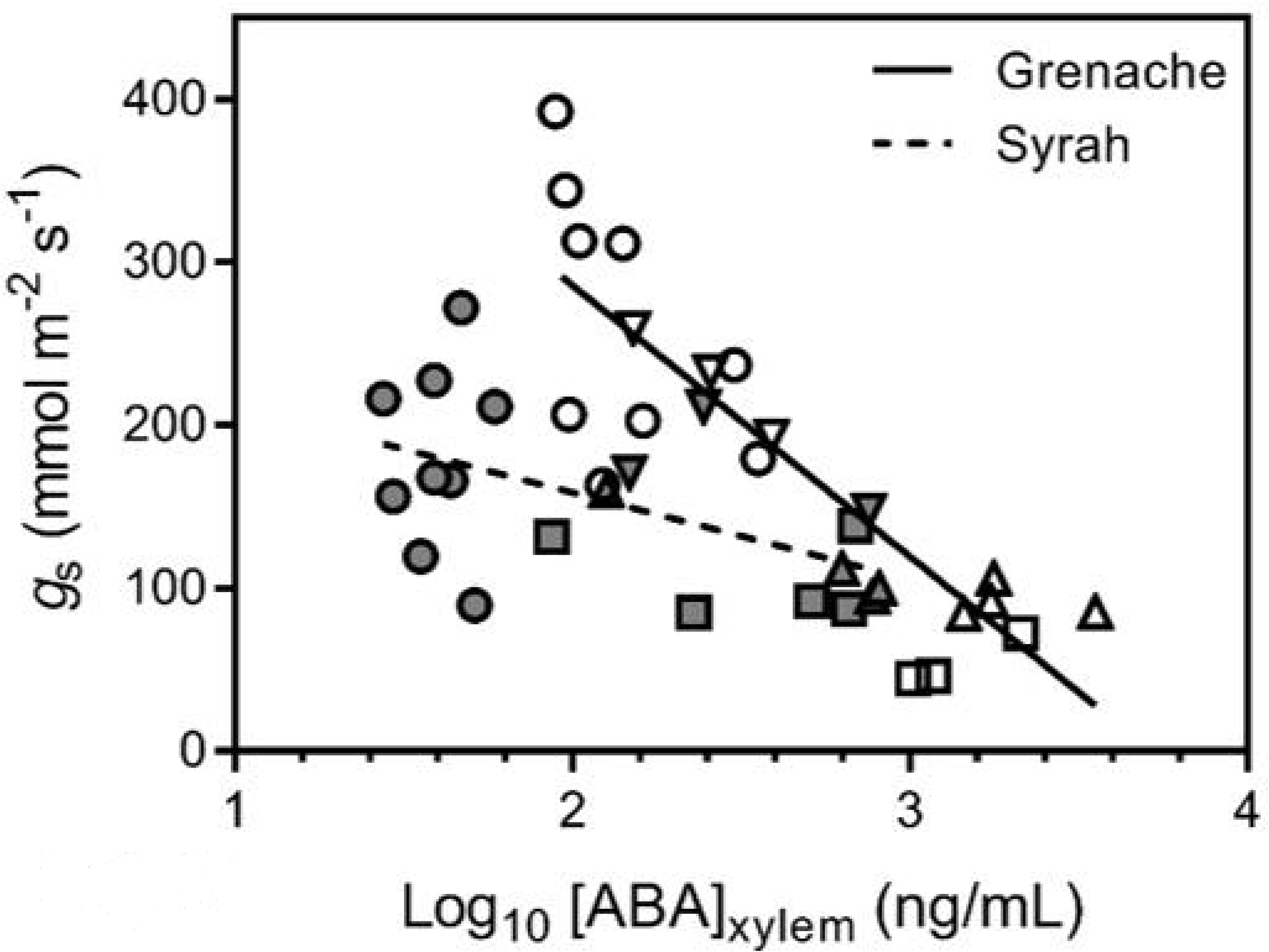
Relationship between [ABA]_xylem_ and *g_s_* measured for well-watered (circles), water deficit (Day 5: triangles up, Day 7: triangles down), and ABA treated vines (squares) from the cultivars Grenache (open symbols) and Syrah (filled symbols). A significant slope (*P* < 0.001) was predicted for Grenache, while the slope for Syrah was not significant (*P* = 0.069). Significant negative regressions were predicted for Grenache (*g_s_* = -166.4 ± 23.9 x log_10_([ABA]_xylem_) + 618.7 ± 63.1; *P* < 0.0001) and Syrah *(gs* = -53.3 ± 18.3 x log_10_([ABA]_xylem_) + 265 ± 40.7; *P* = 0.009).

Simple linear regressions showed that *K*_leaf_ and *K*_plant_ changed similarly in Grenache and Syrah in response to changes in *E* (Fig. 8a,b). ABA treatment (squared symbols) caused a similar response in *K*_leaf_, *K*_plant_, and *E* and, hence, data did fit on the same regression lines. In contrast, linear regressions between *E* and *Lp*_r_ showed a significantly positive response in Grenache, but no response in Syrah (Fig. 8c). Interestingly, *Lp*_r_ remained stable in Syrah despite changes in E. *Lp*_r_ in Syrah was at all times similar to *Lp*_r_ in water deficit or ABA treated vines of Grenache.

**Figure 8.**
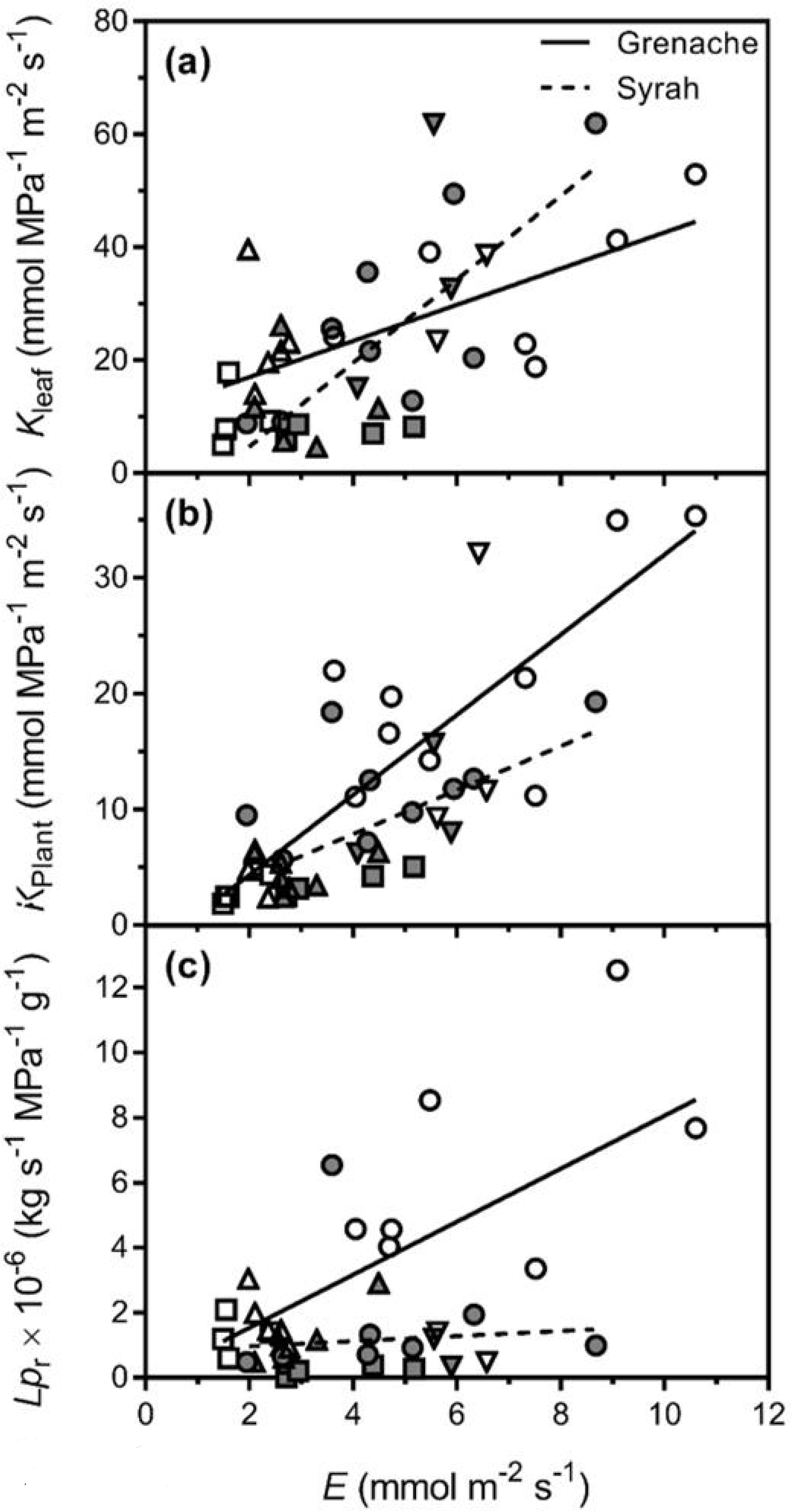
Relationship between *E* and *K*_leaf_, *K*_plant_, or *Lp*_r_ for well-watered (circles), water deficit (Day 5: triangles up, Day 7: triangles down), and ABA treated vines (squares) from the cultivars Grenache (open symbols) and Syrah (filled symbols). (a) *K*_leaf_ was significantly positive correlated to *E* for both Grenache (*K*_leaf_ = 3.2 ± 0.84 x *E* + 10.61 ± 4.41; *P* = 0.0018) and Syrah (*K*_leaf_ = 7.39 ± 1.76 x *E* -10.07 ± 7.98; *P* = 0.0005). The slopes were significantly different (P = 0.0321). (b) *K*_plant_ was positively correlated to *E* for both Grenache (*K*_plant_ = 3.45 ± 0.512 x *E* -2.56 ± 2.66; *P* < 0.0001) and Syrah (*K*_plant_ = 1.899 ± 0.5402 x *E* +0.2659 ± 2.448; *P* = 0.0023) as well. The slopes were not significantly different. (c) *Lp*_r_ was significantly positively correlated to *E* only in the cultivar Grenache (*Lp*_r_ = 0.815 ± 0.2142 x *E* -0.09383 ± 1.081; *P* = 0.0016), but not in Syrah (*P* = 0.6992).

A heatmap of relative gene expression for selected root and leaf AQP genes shows the Log_2_ expression ratio relative to well-watered controls (Fig. 9). Most of the genes measured were down-regulated during water deficit, ABA treatment, and also recovery. The strongest down-regulation was measured for *TIP2;1* in roots of Syrah under WD (Fig. 9b). Only *TIP1;1* was consistently up-regulated in water deficit relative to well-watered vines (Fig. 9). In most cases, gene expression was similarly regulated by water deficit and ABA treatment.

**Figure 9.**
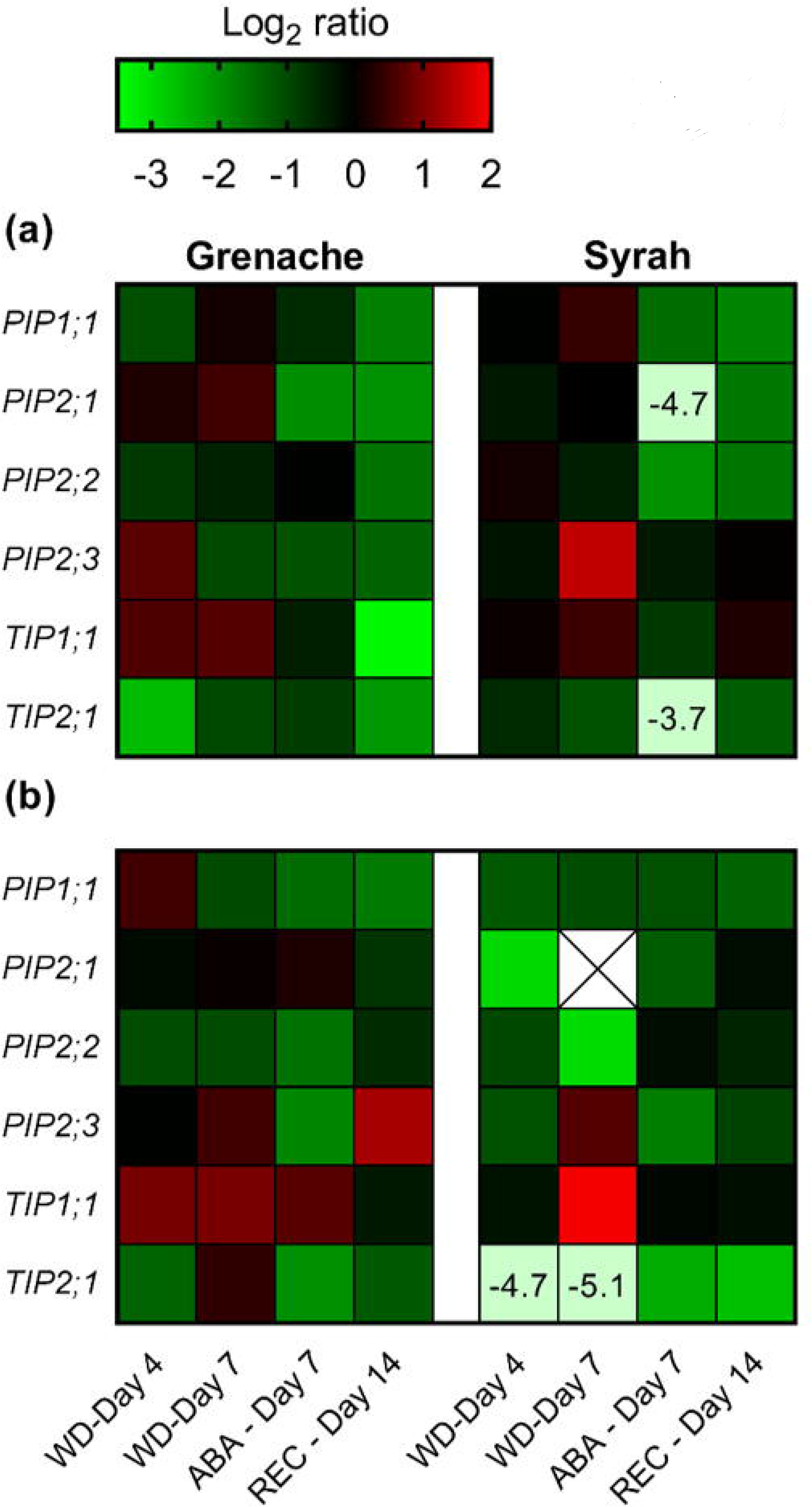
Relative expression of selected aquaporin genes in leaves (a) and roots (b) of water deficit (WD), abscisic acid treated (ABA), and recovery (REC) vines of the cultivars Grenache and Syrah in comparison to well-watered vines. Log_2_ ratios are shown as mean of 2 – 5 biological replicates. Missing data are marked with X. Expression ratios of strongly down-regulated genes are shown as numbers on light green background. The double gradient colour legend shows log_2_ ratios from -3.5 to 2.0.

## DISCUSSION

Isohydric behaviour of plants under water deficit is characterized by a limited decline in Ψ_leaf_ by closure of stomata, while anisohydric behaviour is characterized by maintenance of high *g_s_* and consequently a greater reduction in Ψ_leaf_. There is much interest in these different behaviours since agriculturally important plants with different degrees of isohydry/anisohydry can have very different water demands under certain environmental conditions and hence require different management strategies. The hydraulic and gas exchange properties of plants that confer either isohydry or anisohydry are still relatively poorly understood though there is evidence to indicate that the response of *K*_leaf_ to ABA may be a key feature (Coupel-Ledru *et al.*, 2017). In this study, we investigated additional features of the mechanisms that may account for the differences in isohydry by exploring how both, roots and shoot hydraulics, gas exchange, AQP expression and ABA may be involved. We used two *V. vinifera* cultivars, Grenache and Syrah, which have previously been shown to exemplify these differences (Schultz, 2003). Schultz concluded that Grenache was more isohydric than Syrah as a result of stronger control over *g_s_* and *K*_leaf_. Grapevine has since served as an important model plant to study iso/anisohydric behaviours and several studies have used the cultivars Grenache and Syrah as models for isohydric and anisohydric behaviour, respectively (Coupel-Ledru *et al.*, 2014, Coupel-Ledru *et al.*, 2017, Gerzon *et al.*, 2015, Prieto *et al.*, 2010, Scharwies & Tyerman, 2016, Schultz, 2003, Soar *et al.*, 2006).

### Isohydric and anisohydric behaviour and stomatal sensitivity to ABA

Anisohydric behaviour of Syrah was observed in the present study by the significantly lower Ψ_PD_ and Ψ_leaf_ on Day 7 of the experiment compared to Grenache (Fig. 2e, h), while both cultivars showed a similar relative *g_s_* (Fig. 2b). A higher sensitivity of *g_s_* to changes in Ψ_PD_ for Grenache could also be deduced from the correlogram, where *g_s_* and *E* were positively correlated to Ψ_PD_ only in Grenache. In contrast, *g*_s_ of Syrah was more sensitive to changes in VPD during the mild water deficit experiment as it was observed from the multiple linear regression (Table 1). In the Schultz (2003) experiment, drought stress resulted in a range of Ψ_PD_ comprised of between -0.8 and -1.4 MPa contrasting to the milder and more typical Ψ_PD_ achieved in the present study of -0.3 and -0.5 MPa.

Interestingly, stomata from the cultivar Grenache showed a higher sensitivity to ABA in the xylem of leaves compared to Syrah (Fig. 7). This fits with a higher sensitivity of *K*_leaf_ to ABA fed via the petiole in Grenache compared to Syrah, as observed by Coupel-Ledru *et al.* (2017), although we did not observe a relationship between *K*_leaf_ and [ABA]_xylem_. Tardieu and Simonneau (1998) suggested that isohydric behaviour could be related to stomatal sensitivity to ABA being modulated by Ψ_leaf_ or *E*, while anisohydric behaviour could be due to a single relationship between *g_s_* and ABA. They showed that stomata of maize (isohydric) were more sensitive to ABA at lower Ψ_leaf_. In a leaf dehydration assay on different grapevine genotypes, Hopper *et al.* (2014) found that leaves of Syrah had a higher water loss rate and lower number of stomata per unit area compared to Grenache, which suggested a lower response of stomata to dehydration in Syrah. However, stomata of Syrah showed a quicker response to ABA fed via the petiole to detached leaves compared to Grenache, which was opposite to that expected (Hopper *et al.*, 2014). This may be explained by the effect of Ψ_leaf_ on ABA sensitivity in Grenache. Leaves in the assay would have had a water potential approaching zero, since they would be equilibrating to the feeding solution. Therefore, ABA sensitivity may have been at its lowest in Grenache. Exogenous ABA application in the present study, which was done on intact plants, resulted in the same relationship between *g_s_* and [ABA]_xylem_ compared to ABA responses generated by the mild water deficit (Fig. 7, square symbols). It is important to note that Ψ_leaf_ and Ψ_stem_ were the same with ABA treatment and water deficit for both cultivars (Fig. 2h) indicative of a hydraulic restriction within the plant due to ABA treatment. The down-regulation in response to ABA of several AQPs and the decrease of hydraulic conductances in leaves, root, and whole plant as shown in Fig. 8 (squared symbols) could explain this.

It should also be considered that Grenache may produce more ABA or Syrah could have a higher rate of ABA catabolism. The maximum ABA in the xylem sap of leaves measured during the experiment was higher in Grenache compared to Syrah (Fig. 7). Rossdeutsch *et al.* (2016) found that average ABA concentrations in the xylem sap of Grenache shoots during drought stress were similar to Syrah shoots, however, they measured significantly higher concentrations of dihydrophaseic acid (DPA), a degradation product of ABA, in Syrah compared to Grenache. This may indicate that Syrah has a higher catabolism of ABA, which may regulate its responsiveness to water deficit, and results of the present study support this hypothesis. We observed that water-stressed Syrah vines had nearly three-fold higher DPA concentration in the xylem sap of leaves compared to WW vines as well as all Grenache vines (data not shown). The higher DPA levels suggest greater cumulative water stress in Syrah compared to Grenache as ABA catabolism results in accumulation of DPA and phaseic acid (PA) over time. The higher ABA sensitivity of stomata we observed in Grenache fits with the hypothesis that isohydric behaviour relates to tighter stomatal regulation, possibly through ABA sensitivity. Future research could aim to test the stomatal sensitivity to ABA in Grenache and Syrah at different Ψ_leaf_ to verify the hypothesis by Tardieu and Simonneau (1998).

### VPD effects on stomatal conductance in well-watered vines

As previously observed for several woody and herbaceous species (Franks *et al.*, 2007), including grapevines (Prieto *et al.*, 2010, Schultz, 1996, Soar *et al.*, 2006), positive relationships between *g_s_* and VPD were observed under well-watered conditions for both cultivars. A positive relationship as observed for changes of VPD in the range of 2-3 kPa in this study was also shown by Rogiers *et al.* (2011). For higher values of VPD, a negative effect on *g_s_* should be expected under WD conditions. VPD has been reported to be one of the main sources of variation in *g_s_* in grapevines (Schultz & Stoll, 2010). However, it is not known which mechanisms (passive feedback, active feedback, feedforward, or some combination of these) coordinate *g_s_* and leaf water balance when changes in Ψ_leaf_ are solely due to VPD (Buckley, 2005, Bunce, 1997, Franks *et al.*, 2007, Jones, 1998, Monteith, 1995). The response of *g_s_* to VPD seems to differ between field or laboratory experiments, and between isohydric and anisohydric plants (Novick *et al.*, 2016, Tardieu & Simonneau, 1998). VPD and consequently, *E*, is usually higher in the field (than indoors), where a ‘midday depression’ of *g*_s_ is traditionally attributed to high VPD (Chaves *et al.*, 2010, Correia *et al.*, 1995). Such decrease in *g*_s_ during the afternoon was mostly observed when plants were under mild water deficit, particularly isohydric plants (Bunce, 1997, Tardieu & Simonneau, 1998). In contrast to our observations on Syrah, *g*_s_ of anisohydric species were reported to be similar in the morning and afternoon regardless of VPD suggesting no direct effect of VPD on stomatal control (Tardieu & Simonneau, 1998). Based on a study on the anisohydric cultivar Semillon, Rogiers *et al.* (2012) suggested that stomatal sensitivity to VPD could be higher in dry soils due to the increased levels of ABA. This hypothesis is also supported by results from partial root zone drying in Syrah (Collins *et al.*, 2010). Possibly, VPD effects are dominant in anisohydric cultivars up to a certain point when the soil becomes very dry and stomata then respond more to soil water deficit. Since isohydric cultivars are more sensitive to changes in soil water availability, VPD may have only a significant effect during well-watered conditions. In the present study, we found that both cultivars responded to VPD, but Grenache had a higher *g_s_* at any given VPD under well-watered conditions and the correlation was less strong than that for Shiraz.

Although *g_s_* responds to multiple plant signals triggered in general by the environment, hydraulic conductance of the plant plays a key role in the response of *g_s_* to changes in leaf water relations (Sperry *et al.*, 2002). In our experiment, *K*_leaf_ from both cultivars and *K*_plant_ in Grenache, responded to VPD but no response was observed for *Lp*_r_. These observations agree with the findings by Ocheltree *et al.* (2014) who observed that *K*_leaf_ but not *Lp*_r_ was correlated to changes in *E* caused by changes in VPD. Similarly, *K*_leaf_ in Arabidopsis increased when the plants were transferred to low humidity conditions, and root AQPs were suggested to play a role in the response (Levin *et al.*, 2007). In our experiment, significant positive correlations were found between the gene expression of *PIP1; 1* and *PIP2;2* in roots of Grenache and *E*, VPD, and *K*_plant_ (Fig. 3a). This may indicate that these AQPs had an influence on *K*_plant_ through actions in the root. In Syrah, *K*_leaf_ positively correlated with the same two AQP isoforms in roots, but no correlation was found with *K*_plant_ or *Lp*_r_. As proposed by Simonin *et al.* (2015), changes in *E* are synchronized with changes in *K*_leaf_ to balance the water potential gradient (Ψ_stem_-Ψ_leaf_). While both cultivars in our study showed a similar response of *K*_leaf_ to *E*, only Grenache showed trends to changes in *K*_plant_ and *Lp*_r_ suggesting that this near-isohydric cultivar had tighter control of its internal hydraulic conductances relative to Syrah. In this respect, a higher *Lp*_r_ could minimize the water potential gradient from soil to leaf (Ψ_PD_-Ψ_leaf_) at increased VPD, allowing plants to maintain higher rates of *g*_s_ and preventing a decline in *E* in response to increasing VPD (Pantin *et al.*, 2013). *Lp*_r_ deserves further investigation for its role in the response of *E* at higher values of VPD in both cultivars.

### Divergent strategies for hydraulic control of plant water relations

Control of hydraulic conductivities in roots, stems and leaves is believed to be another important aspect of isohydric and anisohydric behaviour. Schultz (2003) suggested differences in *K*_leaf_ between Grenache and Syrah could be the origin of their isohydric and anisohydric behaviour, respectively. In the present study, *K*_leaf_ was higher in Grenache, which could be explained by its larger xylem vessels (Gerzon *et al.*, 2015), compared to Syrah, however, xylem vessel sizes were not measured in this study. Stronger positive correlations between gas exchange parameters and hydraulic conductances in the near-isohydric cultivar Grenache compared to the anisohydric cultivar Syrah suggest that hydraulic pathways inside isohydric plants are more adaptive to changes in *E* (Fig. 3b). Significant differences in the response of *Lp*_r_ to changes in *E* separated the two cultivars (Fig. 8c), which is an important new finding since little is known about root hydraulics in isohydric and anisohydric cultivars (Lovisolo *et al.*, 2010). Distinct differences in *Lp*_r_ were previously linked to different expression patterns of AQPs hypothesized to result from a xylem-mediated hydraulic signal, possibly from shoots to roots (Vandeleur *et al.*, 2014). The linear regressions between *E* and *Lp*_r_ calculated in this study for Grenache were similar to the linear regressions shown by Vandeleur *et al.* (2009). Interestingly, Coupel-Ledru *et al.* (2017) found that *K*_leaf_ of detached leaves fed with solutions of different ABA concentration decreased with increasing ABA concentration in Grenache, but no change in *K*_leaf_ was observed in Syrah. This response is similar to what we observed in roots (Fig. 8c). While *K*_leaf_ in this study showed a significant positive relationship with *E* for both cultivars (Fig. 8a), no significant correlations were found to [ABA]_xylem_ (Fig. 3b). Different results may be due to the different behaviour of leaves when detached from the plant.

Simonin *et al.* (2015) suggested that a positive correlation between changes in *E* and *K*_leaf_ could help to stabilize the gradient between Ψ_stem_ and Ψ_leaf_ for hydraulic transport to the leaf. This would cause less variation in Ψ_leaf_, which is a feature of isohydric behaviour, and maximise *g*_s_ and, therefore, CO_2_ uptake for photosynthesis. At the whole plant level, our results showed no significant changes in gradient between Ψ_PD_ and Ψ_leaf_ in relation to changes in *E* for both cultivars despite differences in hydraulic regulation, a behaviour termed ‘isohydrodynamic’ by Franks *et al.* (2007). Furthermore, considering the correlation between Ψ_PD_ and Ψ_leaf_ found in our study, both cultivars should be classified as anisohydric as the slope of the relationship between Ψ_PD_ and *E* was not significantly different. However, *g_s_* was differently regulated, as shown by the association between *g_s_* and Ψ_PD_, where Grenache had a steeper slope than Syrah. Given that the root and plant hydraulic conductance in Grenache were closely correlated to *g_s_*, then it is possible to consider an isohydrodynamic behavior in Grenache in which a strong stomatal control maintains relatively constant internal water potential gradients but, at the same time, allows Ψ_leaf_ to fluctuate in synchrony with soil water potential. This new perspective of water transport regulation in plants was recently examined in a study where a theoretical framework, based on the relationship between midday and predawn leaf water potentials, is used to characterize the plant responses to drought (Martínez-Vilalta *et al.*, 2014). The continuum between isohydry and anisohydry requires further exploration yet the functional significance and mechanism of both still remains under debate.

The coordination between *E* and hydraulic conductances could be important for plants to maintain water potential gradients especially when *E* is reduced by low *g_s_*. It is interesting in this respect how ABA application to roots resulted in lower Ψ_stem_ and Ψ_leaf_ despite stomatal closure and no soil water deficit. If *E* is reduced in response to water deficit, e.g. through a reduction of *g*_s_, and hydraulic conductances from the soil to the leaves remain constant, the gradient for water flow (i.e. Ψ_PD_ - Ψ_leaf_) would decrease. Since isohydric grapevine cultivars like Grenache have a more sensitive stomatal regulation to water deficit, the mechanism of decreasing hydraulic conductance with decreasing *E* could be more important. In contrast, decreasing hydraulic conductance in anisohydric plants like Syrah, which maintain a higher *g_s_* and *E*, would cause a stronger drop in Ψ_leaf_.

Aquaporins could play a major role in the regulation of hydraulic conductances. As suggested by Vandeleur *et al.* (2014), AQPs are most likely involved in the regulation of *Lp*_r_ in response to changes in *E*, which was explained by shoot-to-root signaling (Vandeleur *et al.*, 2014). Pou *et al.* (2013) found that in grapevine expression of leaf *TIP2;1* was well-correlated to changes in *g*_s_ during drought and rehydration. In this study, we also found that *TIP2; 1* and *PIP2;2* were positively correlated to *E* and *g*_s_ in roots and leaves of the cultivar Grenache, but not in Syrah (Fig. 3b). The same AQPs were also positively correlated to *K*_plant_, but not to *K*_leaf_ or *Lp*_r_. This difference between Grenache and Syrah could explain different responses in hydraulic conductance to changes in E. It was surprising that no correlation was found between *K*_leaf_ and *Lp*_r_, but there may be other factors that influence this relationship. Clearly, expression of most genes in this study was down-regulated during drought and even ABA treatment (Fig. 9). The down-regulation in response to ABA could explain the decrease of hydraulic conductance in leaves, roots, and whole plant as shown in Fig. 8 (squared symbols). Contrary to our hypothesis, relative gene expression of AQP isoforms in this study was similar between both cultivars (Fig. 9). Therefore, stronger hydraulic control in Grenache could not be explained by stronger regulation of AQP transcripts. However, more significant correlations between AQP gene expression, *E*, and *K*_plant_ in Grenache could indicate that AQPs were more relevant in this cultivar. Overexpression of *SlTIP2;2* in tomato resulted in higher *E* in mutant plants and made them more anisohydric (Sade *et al.*, 2009). The authors suggested that increased membrane permeability for water, i.e. increased hydraulic conductance, could be the cause for the observed increase in *E*. Pantin *et al.* (2013) proposed a hydraulic feed-forward signal for stomatal regulation, which could be mediated by AQPs, as shown by Shatil-Cohen *et al.* (2011). According to their research, AQPs would modulate *K*_leaf_, e.g. via the bundle sheath-mesophyll-continuum, which would send a feed-forward signal to stomata. Hence, two different hypotheses exist for the connection between *E* and hydraulic conductance. Either (i) changes in *E* affect hydraulic conductance through the function of AQPs, or, (ii) a hydraulic feed-forward signal mediated by AQPs affects *g_s_*. In the first case, different stomatal behaviour between isohydric and anisohydric plants could require alternative hydraulic regulation via AQPs. In the second case, alternative hydraulic regulation by AQPs, potentially via ABA, could affect the isohydric and anisohydric behaviour through feed-forward signaling to stomata. Future research could aim to understand which model is more likely.

## Acknowledgments

We would like to thank Trevor Garnett (Plant Accelerator) for support with the DroughtSpotter, and Everard Edwards and Annette Boettcher (CSIRO) for assistance with the ABA analysis. This study was carried out with financial support to Silvina Dayer from the Department of Education of the Australian Government through an Endeavour Research Fellowship and from the Australian Research Council Centre of Excellence in Plant Energy Biology (CE140100008).

